# A novel method for sensor-based quantification of single/multi-cellular traction dynamics and remodeling in 3D matrices

**DOI:** 10.1101/2020.09.24.311647

**Authors:** Bashar Emon, Zhengwei Li, Md Saddam Hossain Joy, Umnia Doha, Farhad Kosari, M Taher A Saif

**Affiliations:** Department of Mechanical Science and Engineering, University of Illinois at Urbana-Champaign, Urbana, IL; Department of Molecular Medicine, Mayo Clinic, Rochester, MN

## Abstract

Cells *in vivo* generate mechanical forces (traction) on surrounding 3D extra cellular matrix (ECM) and cells. Such traction and biochemical cues may remodel the matrix, e.g. increase stiffness, which in turn influences cell functions and forces. This dynamic reciprocity mediates development and tumorigenesis. Currently, there is no method available to directly quantify single cell traction and matrix remodeling in 3D. Here, we introduce a method to fulfil this long-standing need. We developed a high-resolution microfabricated sensor which hosts a 3D cell-ECM tissue formed by self-assembly. It measures cell forces and tissue-stiffness and can apply mechanical stimulation to the tissue. We measured single and multicellular force dynamics of fibroblasts (3T3), human colon (FET) and lung (A549) cancer cells and cancer associated fibroblasts (CAF05) with 1 nN resolution. Single cells show significant force fluctuations in 3D. FET/CAF co-culture system, mimicking cancer tumor microenvironment, increased tissue stiffness by 3 times within 24 hours.

## Introduction

Cell traction is a key mediator of mechanotransduction, which helps cells maintain their size and shape, guide tissue development and homeostasis, support various physiological and pathological processes e.g. wound healing fibrosis ^3^, angiogenesis ^4^, migration ^5,6^, and metastasis ^7–11^. Most importantly, cell contractility creates a link between physical cues and chemical signaling; hence establishes a dynamic reciprocity ^7,12–15^ between cells and the surrounding microenvironment involving neighboring cells and the extra-cellular matrix (ECM). As a result, measuring traction force and assessing downstream effects, such as signaling and matrix remodeling, is very important for understanding numerous biological processes and disease progression. Methods to quantify cell traction on 2D substrates have been developed and advanced over the last 3 decades. However, cells *in vivo* are in 3D environment with extra-cellular matrics (ECM) around them. To date, there is no method to directly quantify cell traction and cell induced matrix remodeling in 3D. This paper closes this gap by developing a novel method for direct measurement of single cell forces and determination of matrix remodeling in 3D ECM over time.

Cell contractility on 2D substrates were first performed using wrinkles produced by cells on silicone rubber films ^16,17^. But wrinkling of thin films is a nonlinear phenomenon and hence, provides a qualitative output. Later, Wang and colleagues developed a technique to quantify cell forces on 2D polyacrylamide (PA) substrates that allowed computation of traction stresses from substrate deformation using computational methods ^18,19^. This technique, referred to as traction force microscopy (TFM), was effective due to PA hydrogel being optically transparent, tunable for mechanical properties, and linear elastic over a wide range of strains (as high as 70%) for constitutive analysis ^18,20^. Subsequent improvements in data collection, analysis, and rendering have made it possible to measure traction stresses with high spatial and temporal resolution ^21–23^. Other techniques have also been developed such as micro-pillar arrays ^24–26^ or Förster resonance energy transfer (FRET) ^27,28^; but these methods depend on soft elastic materials as 2D substrates and are considerably more complex than TFM.

2D cell culture has provided insights into various biological processes; but cellular response and behavior in 3D environments can be very different ^29,30^. Quantifying forces in 3D fibrous scaffolds like collagen is challenging ^31^ due to i) lack of reliable mechanical characterization of the ECM at cellular scale; since macroscopic mechanical properties of collagen is different from local micro-architectural properties at cellular scale, and ii) continuous local remodeling of ECM by cells. Stout and colleagues ^32^ suggested measuring the mean deformation metrics (MDM) of collagen due to force applied by the cells. These MDMs are quantified solely by the 3D displacement field of the cell-surrounding matrices and can indicate overall shape change of the cell (e.g., contractility, mean volume change, and rotation). However, this kinematics-based method cannot provide any information about cell tractions and does not account for the ECM remodeling. Later, Steinwachs and colleagues ^33^ outlined a computational method to measure cell forces in collagen biopolymers using a finite-element approach. They characterized the non-linear mechanical properties of the biopolymers and developed a constitutive equation to compute traction from the 3D deformation field. Although promising, this method is also limited due to difficulties in measuring accurate deformation fields, the assumptions of constitutive equations (stress-strain relations) and computationally expensive analysis. 3D tissue force was measured using micro fabricated pillar structures in ^34^. However, single cell forces or tissue remodeling could not be measured. Consequently, measurements of cellular traction and remodeling in 3D ECM still remain a challenge.

Here, we develop an ultra-sensitive sensor, integrated with a self-assembled three-dimensional tissue construct with a single or a discrete number of cells. The sensor is micro-fabricated from polydimethylsiloxane (PDMS) and can measure cell traction forces circumventing the necessity of constitutive relations in analytically challenging 3D matrices. With a resolution of ~1nN, the sensor is capable of directly quantifying single cell forces in collagen, using force equilibrium laws. In addition, the sensor can be used as an actuator to measure change in ECM stiffness due to remodeling as a function of time, as well as to apply prescribed stretch or compression on the cell-ECM matrix to explore cell response to mechanical deformation in 3D. Hence, the sensing platform offers a range of application for biophysical investigations of cells and tissues. In this paper, we present the details of the sensor and the experimental results that establish novelty, applicability and versatility.

## Results and Discussion

### Concepts and design of the sensor

The basic construction of the sensor comprises of 3 parts-a soft spring, a stiff spring and two grips to hold a self-assembled tissue construct. Fig. 1 A-B presents schematic diagrams and working mechanisms of the sensors. The soft spring (blue spring, Fig. 1A-B) is the force sensing component and the stiff spring (brown spring, Fig. 1A-B) helps to hold the tissue in plane. Their stiffness is denoted by Ks and Kr respectively. Each of the springs is connected to a grip so that any force on the grips can be transferred to the springs. The tissue is formed by dispensing a droplet of liquid cell-ECM (rat-tail collagen I) mixture with low cell density on the grips (Fig. 1A, Suppl. Vid. 1). Cell density of the cell-ECM suspension is so chosen that the droplet contains only a single cell or a discrete number of cells. The liquid cell-ECM mixture fills the gaps of the grid and forms a capillary bridge between them (Suppl. Vid. 1). Collagen polymerizes within about 10 mins and results in a tissue with a single cell (Fig. 1A) or multiple cells (Fig. 1B). The initial length of the tissue is denoted by L_0_. As the cell(s) start to activate, it engages with the collagen, elongates and generates contractile force, F. The force is transferred to the grips and the soft spring extends by *d_c_*, giving cell force, *F = K_s_ ∗ d_c_*. As the stiffness K_r_ is very high, extension of the rigid spring is negligible (*F/K_r_* ≈ 0). Hence, length of the contracted tissue is *L_c_ = L*_0_ − *d_c_*.

**Figure 1.**
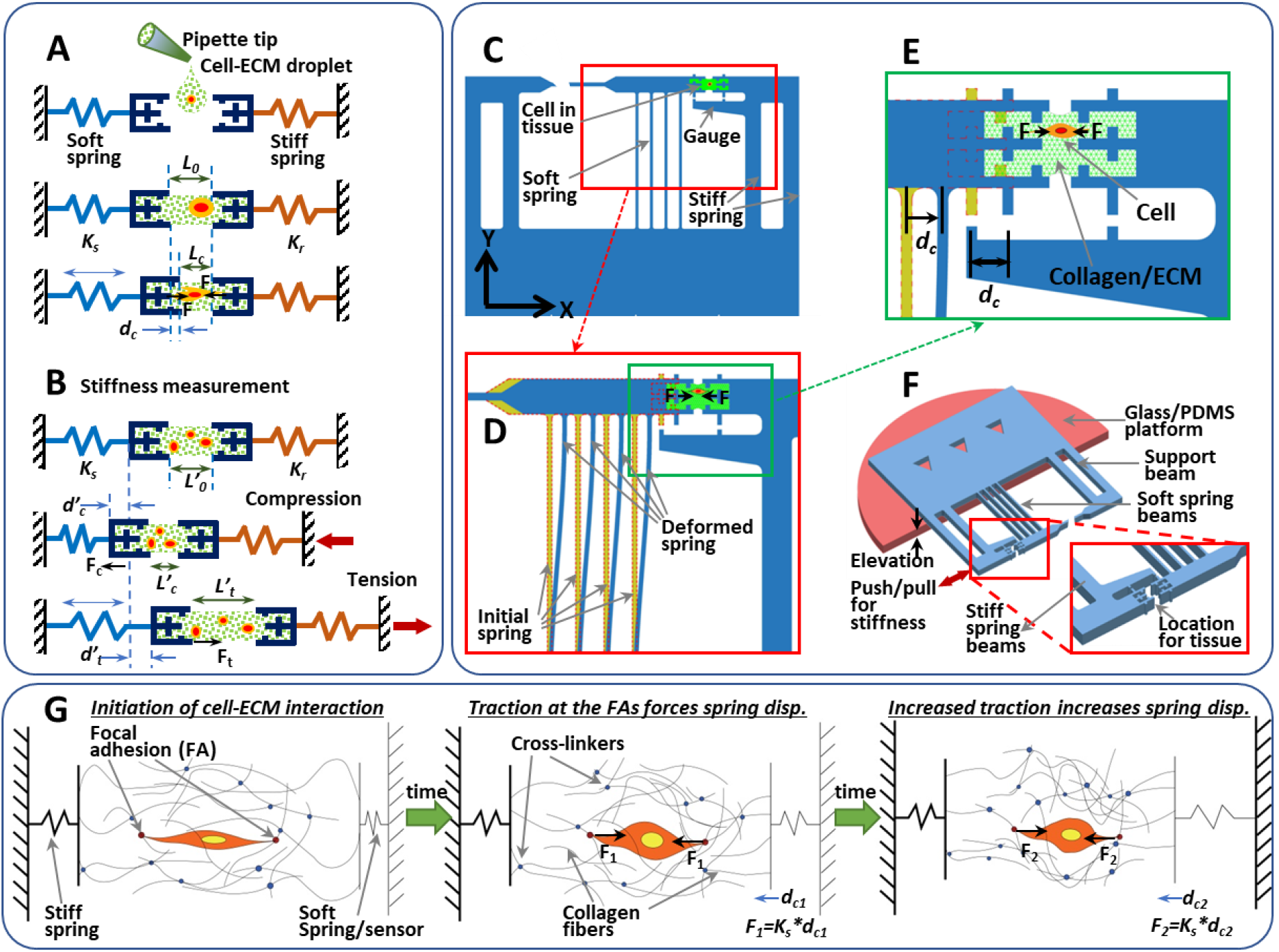
Schematic representation of the concept, design and functional mechanics of the sensor. (A) Simplified model for measurement of cell traction force in 3D matrices. A tissue is formed by dropping cell-ECM mixture between grips connected to springs of known constants. Traction generated by cells in the tissue transfers to the springs and deform the sensing spring (blue). Cell force is quantified as the product of spring constant and deformation (K_s_*d_c_). (B) Technique for measurement of stiffness of the tissue on the sensor. Compressive stiffness can be measured by pushing the stiff spring inside, while tensile stiffness can be measured by pulling the tissue. During the stain application, we continuously monitor the gauges that reads the force as well as strain. (C) Design of the sensor. The thin and wide beams represent the soft and stiff springs respectively. (D) Deformed shape of the beam-springs due to cell traction. (E) Enlarged figure of the tissue and the gauges. (F) Experimental setup (G) Cell activity within 3D ECM and how cell forces transfer through the matric fibers to the springs.

Fig. 1B illustrates the method of measuring the stiffness of the tissue. In order to measure the compressive stiffness of the cell-ECM tissue, the stiff spring is pushed towards the tissue so that it is compressed to a length of L’_c_ from its initial resting length of L’_0_ before actuation. Let d’_c_ be the corresponding deformation of the spring from its rest configuration. Here, the tissue deformation is Δ*L’_c_* = |*L’_c_ − L*’_0_1 and the axial compression force on the tissue (and the springs) is *F’_c_ = K_s_ ∗ d’_c_*. Hence the compressive stiffness of the tissue can be determined as *K_c_ = dF′_c_/d*(Δ*L′_c_*) at Δ*L’_c_*. For measuring tensile stiffness of the tissue, the stiff spring is pulled outwards so that the tissue and the springs are all in tension. And the tensile stiffness of the tissue is determined as *K_t_ = dF′_t_/d*(Δ*L′_t_*) at Δ*L’_t_ = L_t_ − L’*_0_, where F_t_ is the axial tension and deformation is Δ*L’_t_*.

Fig. 1C shows a simple design of the sensor. The thin beams represent the soft spring while the thick beams form the rigid spring. The beams are anchored at one end that allows no rotation or translation. The other ends are rigidly attached to the grid frame that restricts rotation but allows horizontal translation. Fig. 1D shows the deformed beams (spring) when cell(s) apply a force dipole within the tissue. Stiffness of a spring with n beams is *K_s_* = ^12*nEI*^/_*L*^3^_ where E, I and L are modulus of elasticity of PDMS (1.7 MPa), moment of inertia (*I* = ^*hb*^3^^/_12_, with b and h being the width and depth of the beams) and length of the beams respectively. Hence, the stiffness and resolution of the sensor can be controlled by varying the width and length of the beams. For example, *K_s_* = 4.6 *nN*/μ*m*, for one of our fabricated sensors with 4 beams, each 30 μm wide, 200 μm deep, and 2000 μm long. However, it is possible to fabricate thinner beams for sensors with higher sensitivity. An enlarged image of the tissue and gauges for measuring spring deformation is presented in Fig. 1E. A perspective view of the setup is also given in Fig. 1F. It should be noticed that the gauges are placed away from the center of tissue location so that the design allows measurement of force as a function of time without directly illuminating the cell(s). This eliminates the possibility of light induced response of the cells ^54,55^.

Fig. 1G presents a simplified cartoon of force generation by cells and transmission mechanisms to the sensing springs. At first, the cell establishes focal adhesion with the ECM and then starts to contract. As the cell pulls on to the collagen fibers, they get stretched and transmit the tension to the grips, and thus to the sensing springs. During this process, some crosslinks between collagen fibers breaks, while some new ones form. With time, the cell increases contractility that results in higher force output. Again, when the cell retracts from focal adhesions during migration or altering polarities, its traction with the ECM decreases; and the force output also decreases.

### Setup and operation

We employed standard photolithography and deep reactive-ion etching (DRIE) techniques to prepare molds from silicon wafers for the polydimethylsiloxane (PDMS) sensors (details in Methods). Fig. 2A,B show the fabrication process in a setup petri-dish.

**Figure 2.**
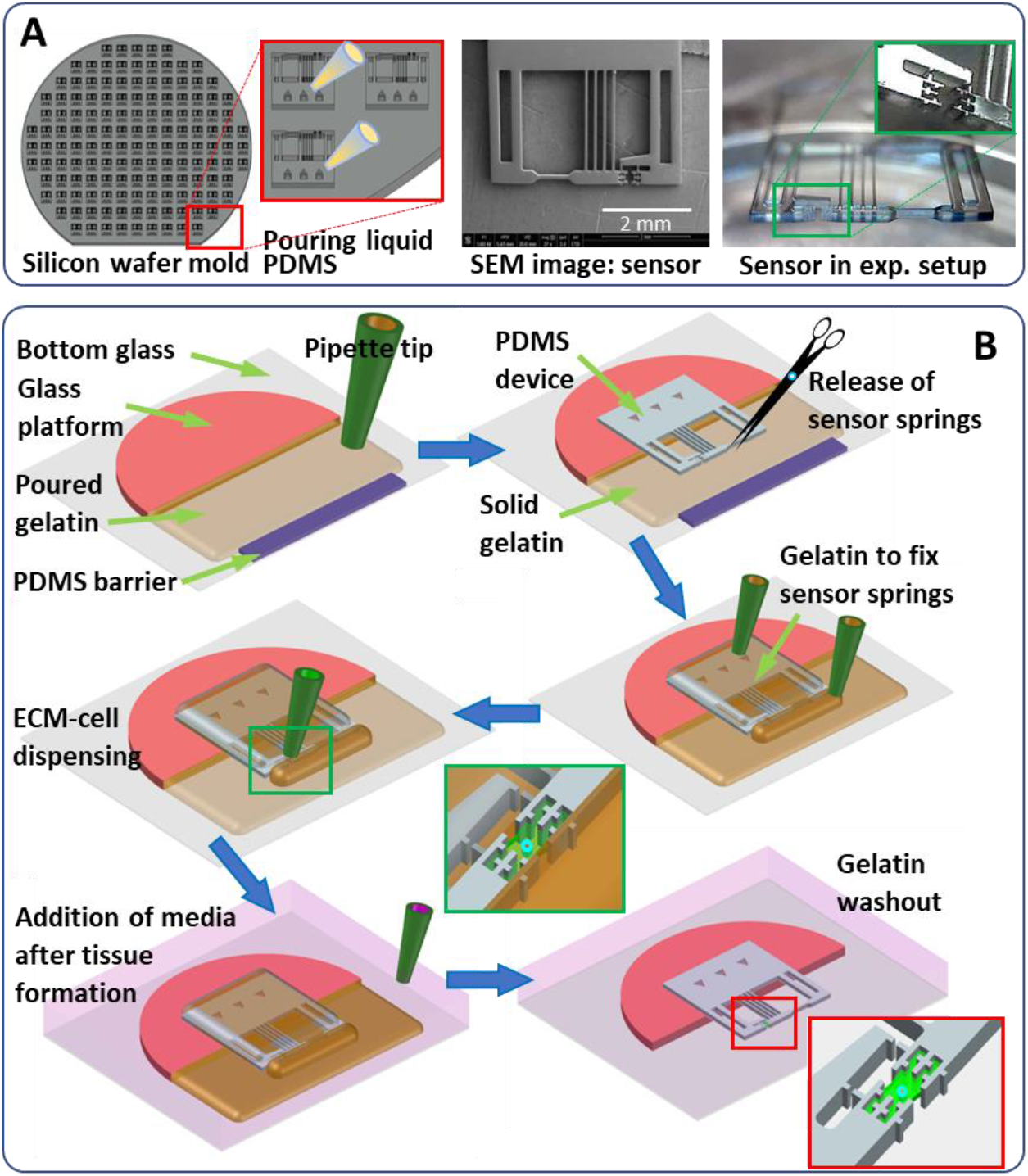
Fabrication of the sensors and preparation of experimental setup. (A) A silicon wafer mold is prepared using microfrabrication. The sensors are cast from the molds by pouring liquid PDMS and curing at 60 °C overnight. SEM image displays a complete sensor. Photo of the setup shows a sensor ready for experiments. The inset presents a zoomed-in image of the tissue grips. (B)Operational scheme to overcome surface energy related challenges for the sensor. Step 1: Sticking glass platform to bottom glass of a petri dish and the device to the platform using liquid PDMS. Step 2: placing a PDMS barrier block underneath the tissue grips. Step 3: Pour liquid gelatin on the sensor beams and parts away from the barrier. Hydrophobicity of PDMS barrier should keep gelatin away from the grips. Step 4: Upon gelation of gelatin at room temp., removal of the barrier to expose the tissue formation site. Step 5: Dispensing of cell-collagen mixture on to the grips to form a tissue bridge. Capillary tension should help collagen to fill out gaps in the grips and later polymerize at RT to form a tissue. Step 6: Submerging the whole setup in cell culture media and incubation at 37° C for 30 mins and then perform washout three times to remove residual gelatin.

In order to have a resolution capable of sensing single cell forces, the sensor springs (beams) must be very soft for high sensitivity. The softness makes the sensor vulnerable to failure due to meniscus forces (surface tension) that appear during its inundation from air to the media. Meniscus forces cause large deformation, buckling and twisting of the sensor beams as well as stiction between them. To overcome this problem, we established an innovative protocol that employs a sacrificial material to protect and anchor the soft beams from exposure to meniscus forces until the ECM is cured and immersed under culture media. We found gelatin to be the ideal sacrificial material for restraining the springs; since it is biocompatible, solid at room temperature and fully dissolves in water at temperatures over 37 degree Celsius ^56^. Fig. 2B shows a step by step process for the experimental setup. At first, an elevated platform is prepared by sticking a glass block to the bottom of a petri-dish. Then a layer of gelatin is placed so that the sensor springs do not stick to the bottom. The PDMS sensor is then attached to the platform using liquid PDMS as glue (Fig. 2B). The connector between the soft spring and its support beam is then severed. Another layer of gelatin is poured on the sensor, only leaving the tissue formation site and the grips unfilled. A droplet of liquid cell-ECM mixture is dispensed between the grips which fills up the space and wets the grips due to capillarity. We then apply a vacuum (~50 kPa, 20 sec) to remove any air bubble. The ECM is then polymerized at room temperature, after which the setup is immersed in cell culture media and incubated at 37.5 °C for about 30 mins. During this period, gelatin melts and dissolves in the media which is later washed out and replaced with fresh media. Thus, the sensor and the tissue bridge never get exposed to air-water meniscus forces. The setup is placed in an environment controlled chamber with microscope for long term imaging of the gauge and the tissue. The gauge gives force readout with time and the tissue images allow monitoring of cellular activities.

### Finite element analysis

In order to investigate how cell position and orientation may affect the force readout from the sensor, we carried out detailed finite element analysis of the sensor-tissue system. PDMS was simulated as a linear elastic material with modulus of elasticity, E = 1.7 MPa ^57^. Collagen was modeled as a bilinear elastic material with compressive modulus significantly lower than tensile modulus (E_tension_ ^1000 Pa^, E_compression_ =0.001 pa) ^33,58,59^. Fig. 3A-D shows the model and results from the finite element analysis (FEA). Three cases were studied where the cell force dipole (two point forces 50 μm apart) was i) aligned ii) at 45° iii) perpendicular to the sensing spring axis (Fig. 3D). The dipole was applied at the center of the tissue and the deformed configuration for the horizontal orientation is shown in Fig. 3B,C. We also varied the dipole force magnitude from 0 to 50 nN. Displacement readout, d_c_ (*F = K_s_ ∗ d_c_*), from the springs shows a linear correlation with applied force (Fig. 3D). For the cell aligned with the spring axis, the sensor reads the total force. With the cell orientation deviating from the spring axis, the sensor reads the total force component along the spring axis.

**Figure 3.**
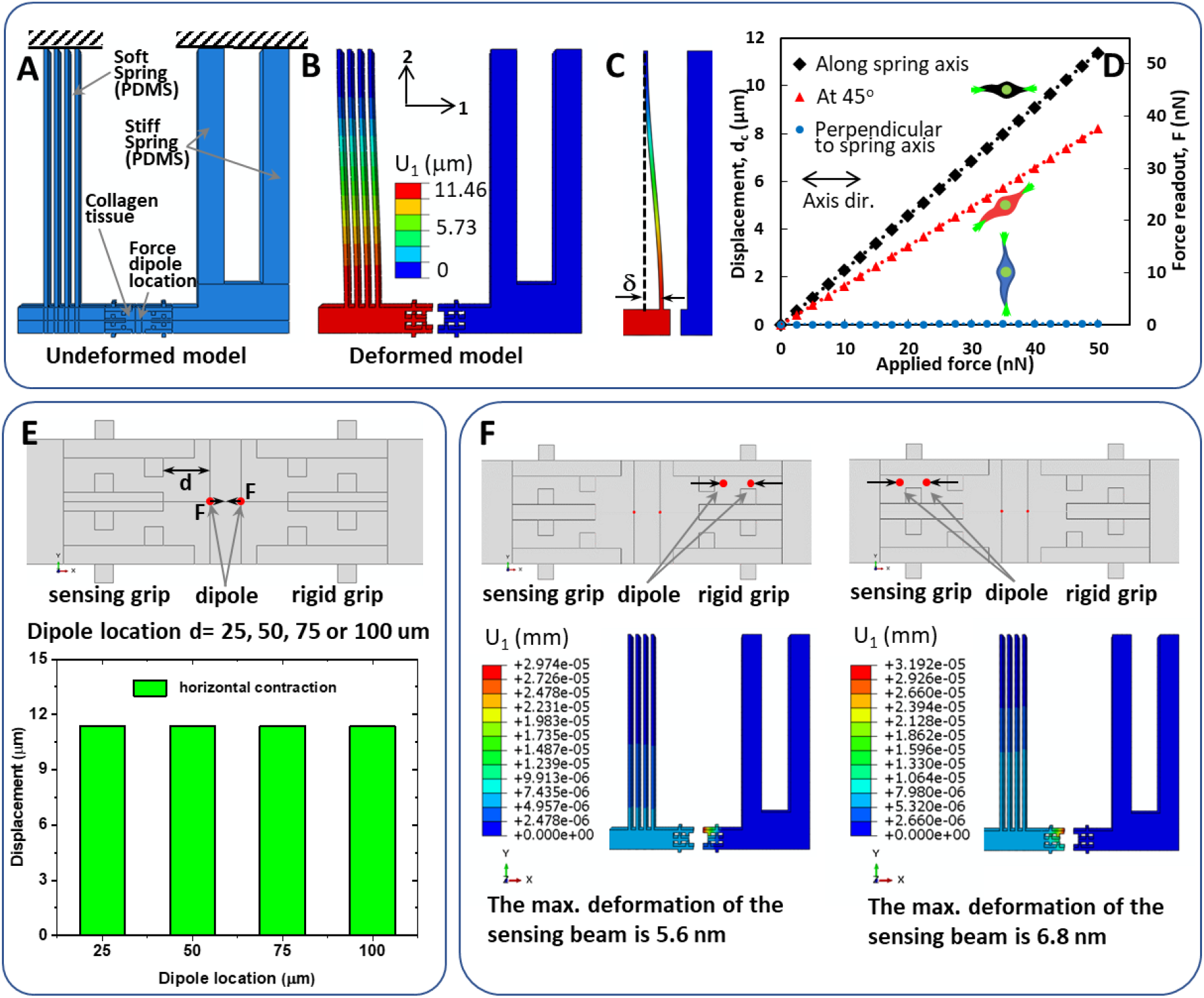
Finite Element Analysis verifies that the sensor can reliably detect cell forces. (A) The undeformed and deformed models show the strains on the beam-springs. The graphs show the relation between cell alignment, force input and the readout from the sensor. Evidently, the sensor precisely detects the force component along the spring axis. (B) Position of the force dipole (or cell force) within the tissue does not affect the sensor’s force detection. (C) Force applied inside the grips of the tissue does not transfer to the sensing spring. Hence, it can be claimed that the force output from the sensor is the result form traction by only the cells residing between the grips.

We also found that the spring deformation, dc, is almost independent of the location of a horizontal force dipole (Fig. 3E). For any position of the applied dipole, readout is within 5% of the applied force. We also found that cells inside the grips do not interfere with the force readout by cells within the tissue (Fig. 3F) In all such intra grip dipole cases, the spring displacement is negligible, ascertaining that the sensor reports data for only those cells that are positioned between the two grips.

### Force and stiffness dynamics

Cell traction and ECM remodeling are continuous inter-dependent processes. We have tested the sensor for time lapse force imaging of single and multiple cells for durations of at least 16 hours. We imaged every 5 minutes for temporal variations in force. However, imaging frequency can be increased without any additional adjustments for experiments that focus on faster response from the cells. Force resolution of the sensor depends on the spring stiffness as well as the image analysis protocol. Our current setup has an image resolution of 167 nm per pixel, which yields a displacement resolution of approx. 17 nm when analyzed with sub-pixel registration in ImageJ ^60,61^. This results in a force resolution of approximately 0.08 nN (K_s_ = 4.6 nN/μm). However, noise analysis with control specimens reveals that the system has a resolution of ~1 nN (Suppl. Fig. 2).

For the current study, we chose NIH/3T3 mouse fibroblasts as a conventional cell line and CAF05 (human colorectal cancer associated fibroblasts) as representative cells for human cancer related experiments. Fig 4A (Suppl. Vid. 2) and 4B (Suppl. Vid. 3) show traction forces generated in the axial direction by a single 3T3 fibroblast and a single CAF05 respectively. Representative phase contrast images of the tissue (cells) are also shown at different time points. We observed that each of these cells initially increase traction gradually, with small fluctuations. During this period, the cells generally probe its microenvironment by protruding small filopodia and pulling the fibers. When the cells elongate and polarize, the magnitude of traction increases and so do the fluctuations. For example, the CAF started to elongate at around 12-15^th^ hours and the force vs time curve also shows spikes in force at the same time. The 3T3 cell exhibits such peaks and drops in forces after 10^th^ hour. These observations are consistent with the fact that cells increase and relax traction periodically while migrating ^21,62^. The maximum force by the 3T3 cells was ~20 nN, while the CAFs generated maximum of ~50 nN force. However, fibroblasts on 2D substrates with stiffness comparable to collagen (~300 Pa) can produce force of ~100 nN ^63^. With higher stiffness, the force can considerably increase to a few micro-newtons. Thus, cell forces measured by 2D TFM can be significantly higher than cell forces in 3D ECM or *in vivo*.

**Figure 4.**
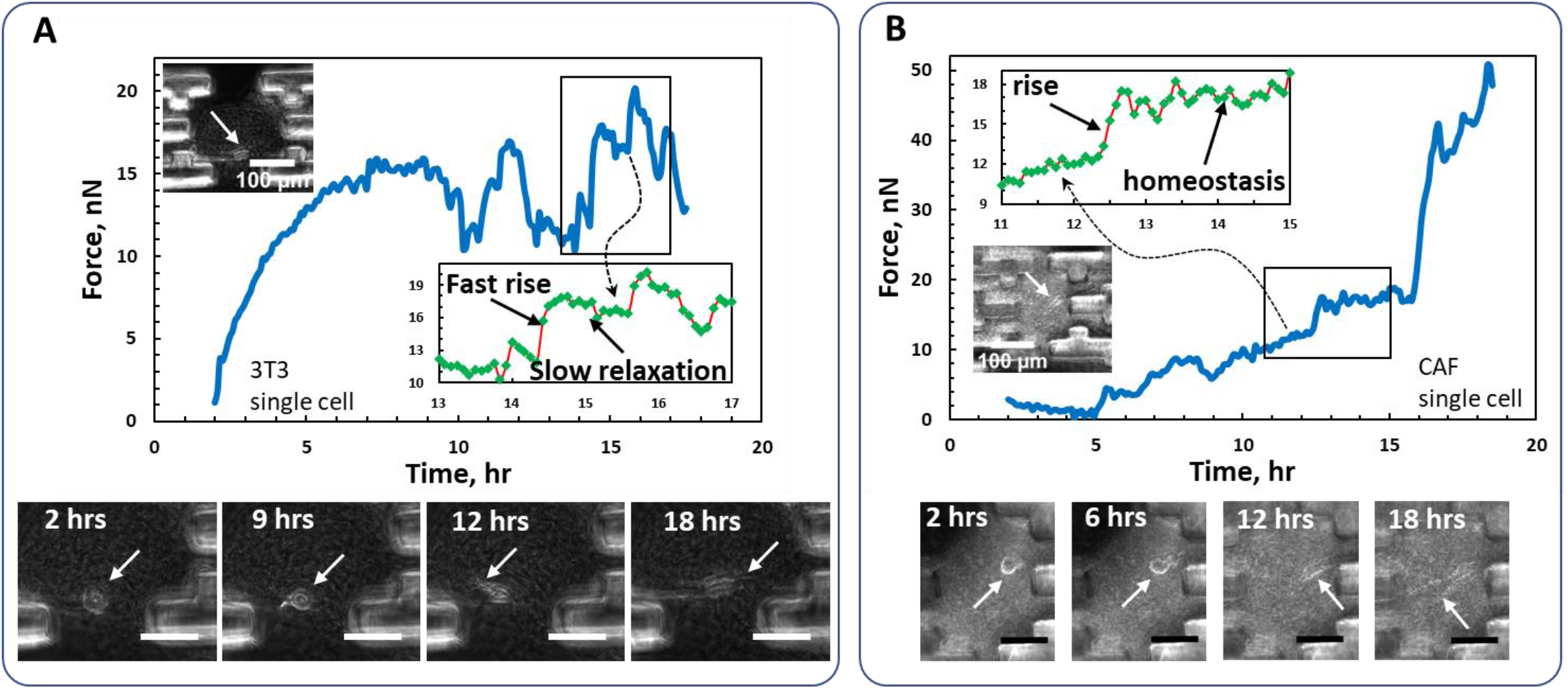
Traction force evolution of single (A) 3T3 and (B) CAF05 fibroblasts. The insets enlarge part of the curves to highlight each data collection point and local dynamics of contraction and relaxation. Phase-contrast images show the cells in the tissue that are generating the force curves. White arrows indicate the location of the cells in the tissues. Scale bars represent 100 μm.

Moreover, the phase contrast images show, for both cases, that when the cells are more elongated i.e. polarized, the magnitude of their forces is much higher compared to the forces when they were not polarized. Insets show finer variations of force. For periods of time, the cells maintain force homeostasis which is evident when the force readout is constant. Another interesting observation is that duration for increasing force is considerably shorter than that for relaxation. This means that the rate of contraction is considerably faster than the rate of relaxation. These findings provide unique insights into cellular traction dynamics in 3D ECM.

We also constructed tissue with a small number of cells to observe the collective traction behavior of fibroblasts. However, the number of cells was kept small enough so that individual cell activities can be tracked. Force evolution with multiple cells demonstrates higher magnitudes, but the magnitude does not increase linearly with cell number (Fig.5A-B). Phase contrast images and corresponding traction force by a collection of 3T3 fibroblasts are presented in Fig. 5A (Suppl. Vid. 4). Images of the tissue show that cells elongate and migrate in the 3D ECM. In addition, similar to the single cell experiments, the curve displays short pulses of traction-relaxation. However, tissues with multiple cells do not demonstrate a faster contraction rate than relaxation, unlike single cells. Data for four active CAFs is shown in Fig. 5B (Suppl. Vid. 5). Similar to single cells, this curve also exhibits periods of steady state, with occasional increase of force. To confirm that the output force is generated by the cells, we applied Y-27632 that inhibits ROCK signaling pathways to relax cell traction. As expected, within 20 minutes, the drug reduced the force by about 60%. After washout at this point, the cells immediately started to contract and re-established pre-drug level of force in about 1 hour. The cells continued to increase force for the following 2 hours. This indicates that the forces detected by the sensors are generated by the cells within the 3D collagen tissue.

**Figure 5.**
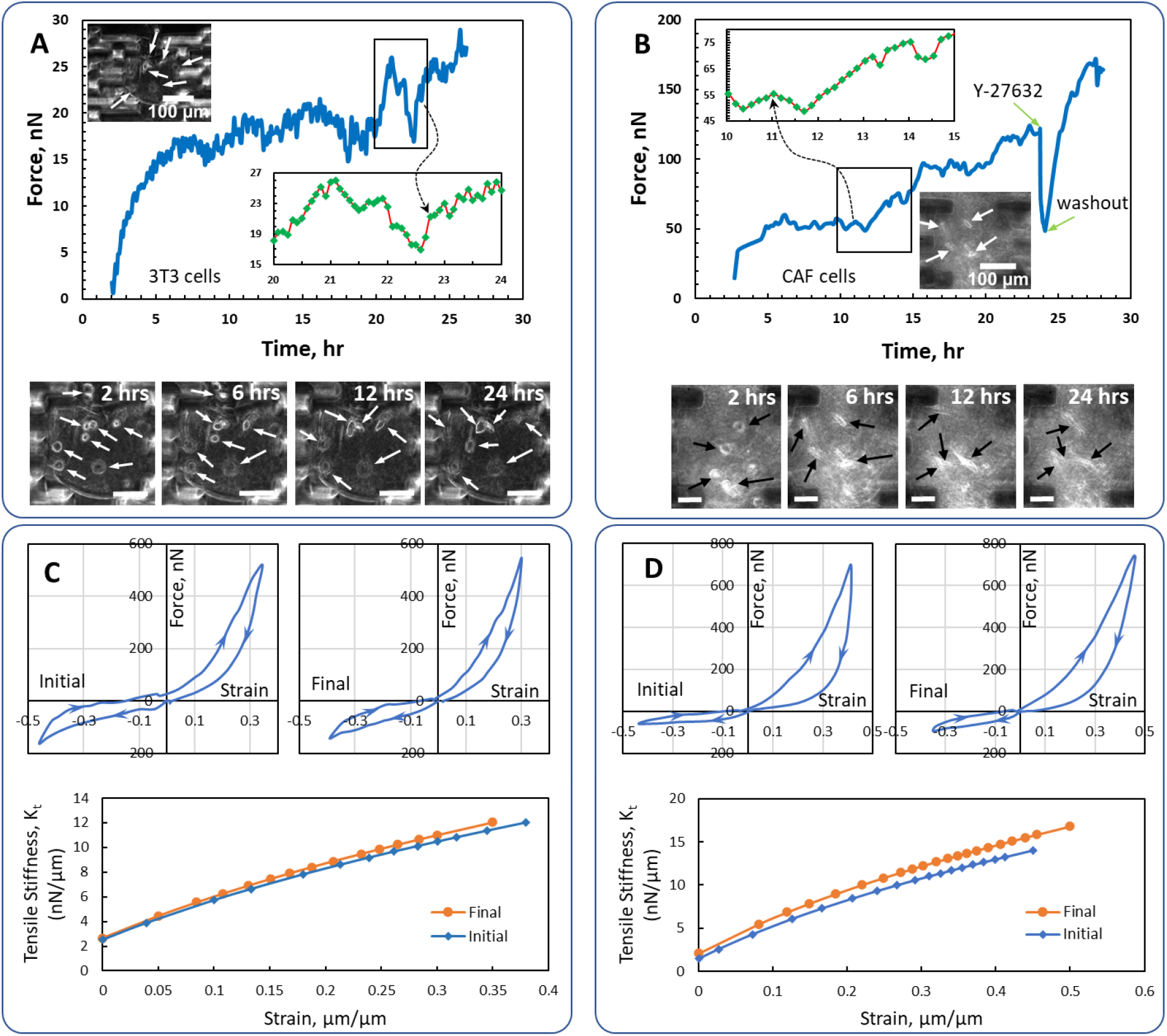
Traction force evolution of multiple (A) 3T3 and (B) CAF05 fibroblasts. The insets enlarge part of the curves to highlight each data collection point and local dynamics of contraction and relaxation. Phase-contrast images show the cells in the tissue that are generating the force curves. Arrows indicate the location of the cells in the tissues. Scale bars represent 100 μm. Force vs. strain curves and corresponding tensile stiffness graphs of the tissues with a number of (C) 3T3 and (D) CAF05 fibroblasts. The compression-tension testing were performed at the start of the experiment when the cells were still not activated and after about 24 hrs. The curves show that the cells increased the stiffness of the tissues very slightly in this period.

Moreover, we assessed remodeling of collagen by the CAFs and 3T3 fibroblasts, in terms of stiffness of the tissue. To this end, the tissue stiffness was measured at the 2^nd^ hour (initial) and 28^th^ hour (final) of tissue formation. Suppl. Vid. 6 shows the process of applying strains on the tissue for stiffness measurement. Force vs strain curves for both compression and tension with 3T3s and CAFs are shown in Fig. 5C and 5D respectively. During the loading-unloading cycle, we performed compression first and then tension. Both compression and tension curves show nonlinear relations. It is also evident that the tissues have higher stiffness in tension compared to that in compression. This indicates that the sensor is capable of reliable mechanical testing of micron scale soft specimens since is can capture behavior of polymeric scaffolds like collagen ^64–66^. The tension-loading portions of the curves were fitted to Mooney-Rivlin model for hyperelastic materials. The tangential tensile stiffness (Kt) was derived from the fitting model and variations of Kt with increasing strains are also presented in Fig. 5C-D (details in Methods and Supplementary materials). Both CAFs and 3T3 cells increased the stiffness of the tissues very slightly in about 24 hours. We understand that the tissue requires more cells or time to make significant remodeling, but the sensor can detect subtle changes in the stiffness.

### Tissue with cancer cells

To demonstrate potential application of the sensor to study in vitro tissues that mimic tumor microenvironment, we created lung cancer and colorectal cancer (CRC) models on the sensors with A549 (human lung epithelial carcinoma) and human CRC cell lines. Fig. 6 demonstrates the sensor’s capability to host a 3D *in vitro* tissue model where cancer and stromal cells reside in close proximity of each other. The confocal images show that the model consists of one FET CRC cluster with 3 cells, five CAF05 stromal fibroblasts and collagen as ECM scaffold. Confocal z-stack images of labeled F-actin and nuclei (Fig. 6A-B) of the cells were used to reconstruct the 3D structure of the cells; while the two-photon second harmonic generation (SHG) images (Fig. 6C) show the collagen organization around the cells. Three-dimensional reconstruction of the tumor tissue (Fig. 6D-J, Suppl. Vid. 7) validates that the model offers a 3D culture condition necessary for simulating tumor microenvironment.

**Figure 6.**
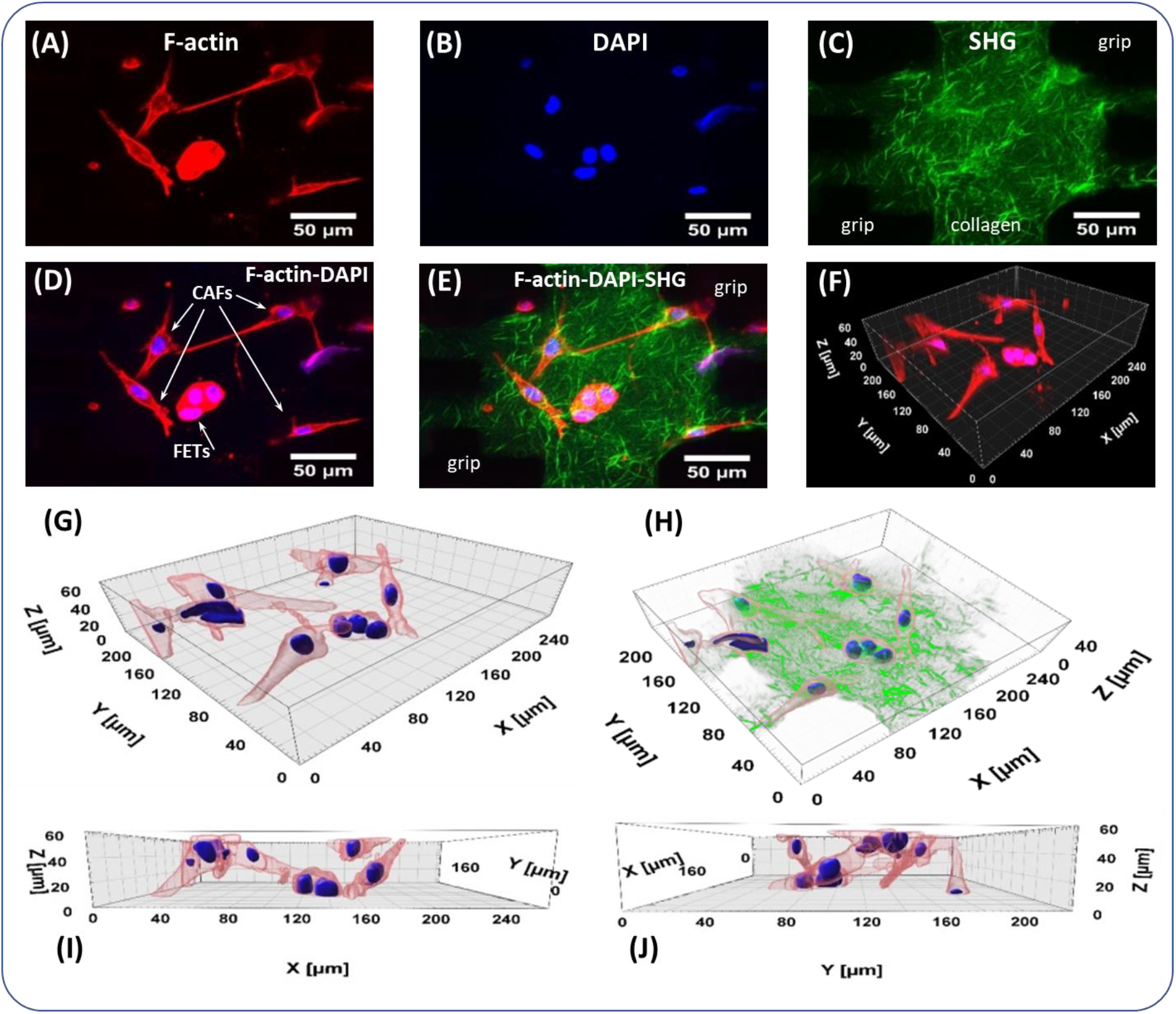
3D tumor model on the sensor with cancer cell and CAF co-culture. (A) Maximum intensity projection of F-actin labeled with phalloidin conjugated with Alexa Fluor 647; (B) Maximum intensity projection of cell nuclei labeled with DAPI; (C) SHG image of collagen fibers; (D) Overlay image of the F-actin and nuclei of the cells; (E) Overlay image of the F-actin, collagen fibers and nucleus of the cells; (F) 3D reconstruction of confocal z-stacks of F-actin and nucleus of the cells; (G) 3D surface rendering of confocal z-stacks of F-actin and nucleus of the cells; (H) overlay image of collagen fibers and 3D surface rendered image of F-actin and cell nucleus; (I) XZ and (J) YZ plane of 3D rendered surface from (G).

For force dynamics evaluation, we constructed three different tissues with (i) A549 cells (Fig. 7A; Suppl. Vid. 8), (ii) FET (human colorectal carcinoma) cells alone (Fig. 7B,D; Suppl. Vid. 9), (iii) FET and CAF05 (human colorectal CAFs) cells together (Fig. 7C,E; Suppl. Vid. 10). Most of the A549 lung cancer cells are initially singular (Fig. 7A, 2 hrs) like the fibroblasts; but with time, they coalesce into a number of small clusters (Fig. 7A, 18 hrs). However, FET cancer cells are different from the fibroblasts in terms of cell-cell adhesion and migration. Initially, FET cells form small clusters (Fig 7B), whereas fibroblasts remain as individual cells (Fig. 5A,B). With time, these small clusters of FET cells combine into larger clusters forming cancer spheroids. The spheroids inside 3D collagen are dynamic; they continuously evolve into different shapes and sizes and also maintain traction with the ECM. However, unlike the FET cells, A549 cells did not form large spheroids, even after 30 hrs.

**Figure 7.**
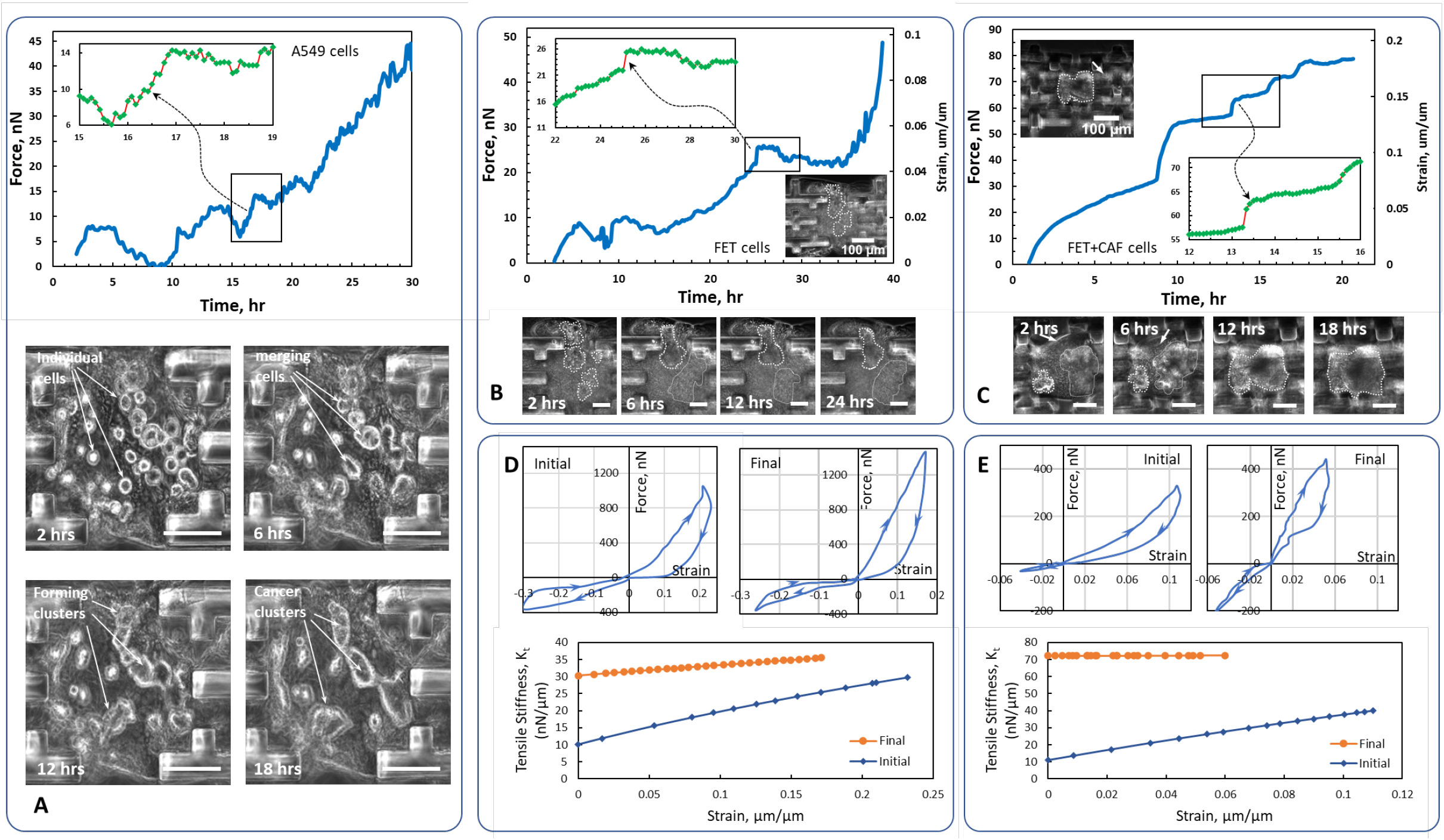
Cell traction and matrix remodeling in cancer. Traction force evolution of a cluster of (A) A549 lung cancer cells (B) FET CRC cells and (C) FET and CAF05 fibroblasts co-culture tissue. The insets enlarge part of the curves to highlight each data collection point and local dynamics of contraction and relaxation. Phase-contrast images show the cells merge into clusters and spheroids in the tissue. White dotted lines indicate cell cluster boundaries, and the arrow indicates the location of a CAF cell in the tissues. Scale bars represent 100 μm. Force vs. strain curves and corresponding tensile stiffness graphs of the tissues for (D) FET only and (E) FET and CAF05 co-culture. The compression-tension testing were performed at the start of the experiment and after about 24 hrs. Both curves show that the cells significantly increased the stiffness of the tissues which indicate substantial remodeling.

**Figure 8.**
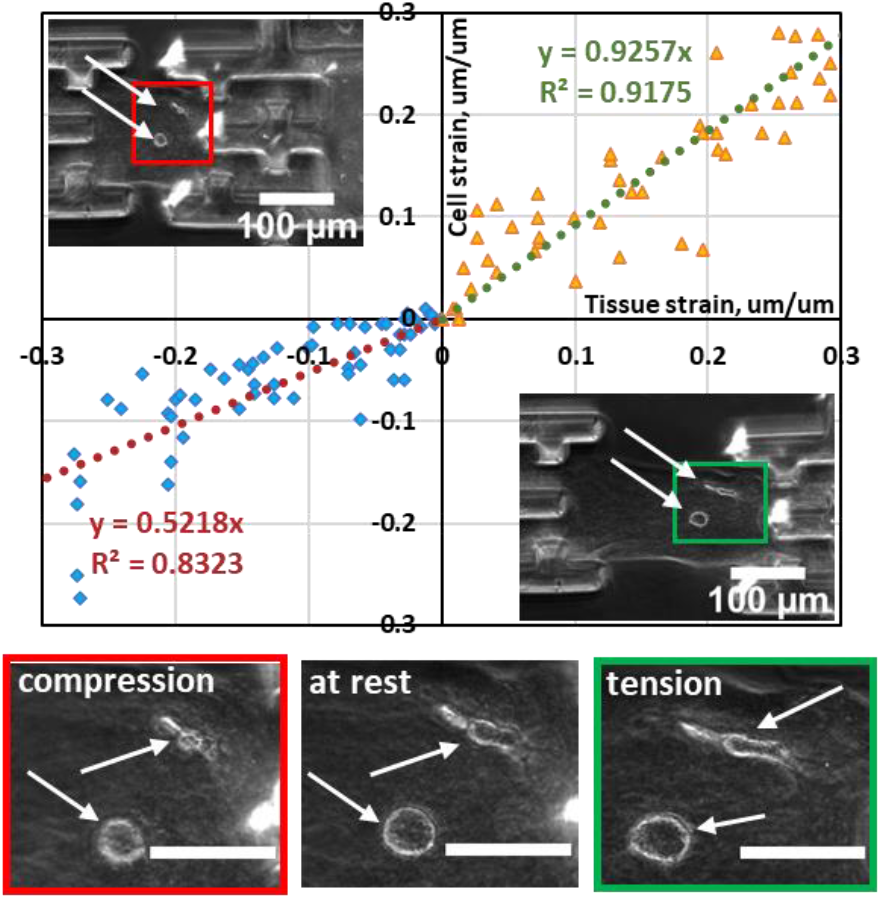
Cell stretch-contraction test results on the sensor. The orange and blue markers represent tension and compression data points respectively. Linear regression lines for the data shows the relationship between applied tissue strain and corresponding cell strain. The phase contrast images and the insets show the cells at different stages of the experiment-compressed, at rest and elongated. White arrows direct to the cells. Scale bars: 100 μm.

Evolution of force within a 3D collagen matrix by A549 and FET cancer cells are shown in Fig. 7A-B. The force history of A549 cells appears to be similar to those of fibroblasts (Fig. 5A). The FET force history (Fig. 7B), however, exhibits a distinct feature, i.e., there are very few drops in force, unlike those for the A549s (Fig. 7A) and fibroblasts (Fig. 5A-B). One potential reason is that the spheroids do not migrate as much as the A549s and fibroblasts (Suppl. Vid. 9). The dynamic force that these spheroids generate, results in significant remodeling of the surrounding ECM. Fig. 7D shows that the cancer spheroids significantly increased (min. 40%, max.200%) the stiffness Kt of the tissue in about 40 hours.

In order to mimic a 3D tumor microenvironment, we created a tumor tissue on the sensor with FETs and CAFs in collagen. Co-culture of cancer and stromal cells facilitates various signaling and cross-talk between them ^7,67,68^. For this specimen, the small FET clusters agglomerate and form a larger spheroid, but the CAFs remain as isolated cells and move around the spheroid (Suppl. vid. 10). Overall, the co-culture specimen showed a stronger force output and ECM remodeling. The force curve in Fig. 7C shows that the tissue slowly increases force without force relaxation at any point and occasionally generate traction at high rates. In addition, this specimen underwent a greater remodeling as indicated by almost three-fold increase in Kt (min. 133%, max. 600%, Fig. 7E). Thus, these experiments demonstrate that the sensor can be a suitable tool for biophysical investigation of cancer cells, stromal cells and the tumor microenvironment (TME). For instance, we can examine the effect of stiffness of TME on cross-talk between cancer and stromal cells, or metastatic migration of invasive cells. It is possible to create *ex-vivo* tumor tissues on the sensors using primary cells derived from biopsy samples and utilizing traction force/stiffness for personalized drug screening.

### Application of strain for cells in 3D ECM

Utilizing the sensor, we applied stretch and compression to the tissue between the grids by applying a prescribed motion on the supporting spring using a piezo stage. The ECM is thus subjected to tensile and compressive strains, which in turn transfers the strains to the cell. Fig. 7 (Suppl. Vid. 6) presents phase contrast images of cells under stretch and contraction, and the relationship between cell and ECM deformation. Cell strain and ECM strain is quantified from the following –

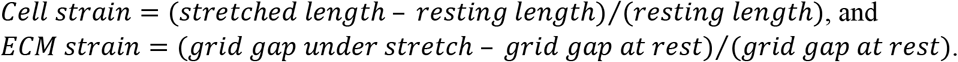

The data suggests that the cell strain is about 93% of the ECM strain in tension; while in case of compression, only 52% of the tissue strain is transferred to the cells. One explanation is that the cell gets stretched by the collagen due to cell-ECM adhesion. Under compression, the fibers buckle and fail to transfer force and strain to the cells.

### Conclusion

We developed a high-resolution sensor that allows self-assembly and culture of 3D tissue models; and described the basic principles of the design and analysis, and its methods of operation. The sensor can report single and multiple cell forces in 3D ECM over a long period of time with a resolution of 1 nN, as well as quantify the change of stiffness of the tissue remodeled by the cells. Feasibility of the sensor was tested by forming tissues with a single 3T3 fibroblasts and CAF05 cells (human colon cancer associated fibroblasts), as well as with multiple cells (3T3, CAF05, human colon cancer cell FET, human lung cancer cell A549). Single 3T3 cells produced a maximum of ~20 nN, while single CAFs produced up to ~50 nN. Their corresponding forces on 2D substrates with similar stiffness approach ~100 nN. Multiple cells exhibit higher overall force collectively, although the magnitude does not scale linearly with the number of cells. In terms of remodeling, 3T3 fibroblasts or CAFs (<10 cells) did not induce any significant change in stiffness of the tissues. On the other hand, FET and A549 cancer cells form clusters, generate large force and remodel the matrix and increase the stiffness by about 2 times in 24 hrs. FET and CAF05 co-culture changes stiffness by about 3 times. Finally, we show that the sensor can be used as an actuator to apply stretch and contraction on the cells and ECM in 3D. In summary, the micro-sensor allows to measure single and multiple cell traction in 3D matrices, to quantify tissue stiffness and remodeling. An array of such sensors can be applied to form tumor environments from patients’ cell for drug screening and prognosis.

## Materials and Methods

### Cell culture

Human primary colorectal tumor CAFs, CAF05 (Neuromics, Edina, MN, USA) were maintained in Vitroplus III, Low Serum, Complete medium (Neuromics, Edina, MN, USA). NIH 3T3 cells, obtained from ATCC (Manassas, VA, USA), were cultured in fibroblast media prepared with Dulbecco’s Modified Eagles Medium (DMEM, Corning), 10% Fetal Bovine Serum (FBS, Gibco) and 1% Penicillin-Streptomycin (Lonza). FET human colorectal carcinoma cells were a gift from the lab of Prof. Barbara Jung, Dept. of Medicine, UIC and were maintained in 89% DMEM/F12 50:50 (Gibco), 10% Fetal Bovine Serum (FBS, Gibco) and 1% Penicillin-Streptomycin (Lonza). A549 lung cancer cells were collected from Prof. Kosari, Mayo Clinic, Rochester, MN. These cells were cultured in F-12K medium (Kaighn’s Modification of Ham’s F-12 Medium, ATCC, Manassas, VA) supplemented with 10% FBS (Gibco). Cells were grown at 37 °C in a humidified incubator with 5% CO_2_.

### Fabrication and assembly of PDMS sensors

The sensors were cast from microfabricated silicon molds. Standard 500 μm silicon wafers (University wafer, Boston, MA, USA) were patterned by photolithography and etched to a nominal depth of 200 μm using the deep reactive-ion etching (DRIE) process (STS Pegasus ICP). Next, the etched wafers were coated with polytetrafluoroethylene (PTFE) to facilitate removal of PDMS from the mold. PDMS (Sylgard 184) base and cross-linker were mixed at 10:1 ratio by weight, pipetted into the molds, and allowed to fill all the features and trenches by capillary mircomolding ^69^. The specimens were cured at 60 °C for 12hours and lifted off the silicon molds. For assembling the setup, a rectangular glass piece (#2 cover glass, Corning) was glued to the bottom glass of a glass-bottom petri-dish (60 mm diameter, Corning) using uncured PDMS. The glass piece served as the elevated platform. The sensors were also fixed to the glass platform using uncured PDMS. We arranged 10-15 single sensors in one 60 mm petri-dish.

### Tissue formation

For preparing all tissue precursor solutions, an ECM solution was prepared on ice by first neutralizing rat-tail collagen I (Corning) with 1N sodium hydroxide, 10X PBS, and deionized (DI) water. We followed Corning recommended protocol ^70^ to prepare a final collagen solution of 2 mg/ml with pH 7.2 from a high concentration stock solution of 8.9 mg/ml in 0.02 N acetic acid. For a single cell in the tissue, cells were suspended in the ECM solution at a density of 150 x10^3^ cells/ml. Cell density was increased linearly based on the desired number of cells in the final tissue construct. Cell-ECM mixture was then pipetted onto the grips of the sensors and was allowed to fill the channels. A syringe pump (NE-1000; New Era, Farmingdale, NY) was employed to control pumping of the liquid mixture through a flexible tube to a fine needle with precise volume and flow rate. The needle was fixed to a 3D automated stage equipped with piezo-actuators with fine steps (few nms) to precisely dispense cell-ECM mixture on to the space between the two grids (Suppl. Vid. 1). All the components are kept at 0°C before dispensing to avoid early polymerization of ECM.

Generally, capillary tension is enough to draw the mixture in and drive the air pockets out; however, removal of persistent bubbles can be aided by applying low-pressure for about 20 seconds in a vacuum desiccator. Until this point, the procedures were carried out on ice in order to delay the onset of polymerization. After removing any remaining air bubbles, the cell-ECM mixture was allowed to polymerize at room temperature for 10-15 mins; then the sensors with assembled tissues were inundated in culture media and placed in the incubator.

### Immunohistochemistry and imaging

The setup was placed, for the duration of the experiment, in an environment controlled chamber enclosing an inverted optical microscope (Olympus IX81, 40X objective, Olympus America Inc., Center Valley, PA) mounted on a vibration isolation table (Newport Corporation, Irvine, CA). The chamber maintains cell culture conditions at 37 °C temperature, 5% CO_2_ and 70% humidity. Moreover, a motorized stage (Prior Scientific Inc., Rockland, MA) allowed automatic imaging of multiple specimens at preset locations at multiple time points. Images of both the tissues and the gauges were acquired in phase contrast or brightfield mode with a Neo sCMOS camera (active pixels 1392 × 1040, resolution of 167 nm per pixel) (Andor Technology, Belfast, Northern Ireland). For calculating spring displacements, images of the sensor gauges were analyzed using template matching plugin in ImageJ with sub-pixel resolution; and the resolution was approx. 17 nm.

For confocal imaging, the samples were fixed with 4% Paraformaldehyde (PFA) in Phosphate-buffered saline (PBS) for 1 hour. Subsequently, 0.2% Triton X-100 in PBS was used to permeabilize the samples and 2.5% Bovine serum Albumin (BSA) with 2% Normal goat serum (NGS) in PBS was used as a blocking solution. Samples were then incubated overnight in Phalloidin conjugated with Alexa Fluor 647 (1:40) (Invitrogen, Carlsbad, CA, USA) at 4°C. Afterwards, the samples were washed with PBS for three times, then incubated in 4’,6-diamidino-2-phenylindole (DAPI) (1:1000) (Invitrogen, Carlsbad, CA, USA) for 10 minutes and washed with PBS again. The image acquisition was done with a confocal microscope, LSM710, using an EC Plan-Neofluar 20X/0.5 NA objective lens (Carl Zeiss AG, Oberkochen, Germany). SHG images are acquired subsequently using the same LSM 710 two-photon excitation microscope and 20X objective. Maximum intensity projection of the acquired confocal z-stacks was constructed using the ImageJ (U.S. National Institutes of Health, Bethesda, Maryland, USA) software and IMARIS (version 9.6.0, Bitplane AG, Zurich, Switzerland) is was used for the 3D surface rendering.

### Finite Element Analysis

Three-dimensional finite element analysis (FEA) was performed using commercial software Abaqus to investigate the beam deformation of the force sensor under the cell contraction. Eight-node brick solid elements (C3D8R) were used to discretize the geometry of force sensor and extracellular matrix, and refined meshes were adopted to ensure the accuracy. The force sensor has four parallel aligned beams with 30 μm gaps in between, and each of the beams has the length of 2000 μm, width 30 μm and depth 200 μm. The cell contraction was simulated by applying a 50 nN dipole forces with 50 μm distance. Linear elastic model was used to demonstrate the material behavior of PDMS force sensor and ECM. The elastic moduli (E) and Poisson’s ratios (ν) used are E=1.7 MPa and ν =0.48 for the PDMS sensor. For ECM, the tensile elastic modulus and compressive modulus are 1 kPa and 0.001 Pa respectively, and Poisson’s ratio is 0.48.

## Supporting information

Suppl. Vid. 1

Suppl. Vid. 2

Suppl. Vid. 3

Suppl. Vid. 4

Suppl. Vid. 5

Suppl. Vid. 6

Suppl. Vid. 7

Suppl. Vid. 8

Suppl. Vid. 9

Suppl. Vid. 10

## Acknowledgements

Research reported in this publication was partially supported by the National Institute of Biomedical Imaging and Bioengineering of the National Institutes of Health under Award Number T32EB019944 to BE. The content is solely the responsibility of the authors and does not necessarily represent the official views of the National Institutes of Health. Funding was also provided by National Science Foundation grant NSF ECCS 19-34991 and Mayo clinic grant PO 66236006. We also thank Dr. Lin Yang for assistance with the A549 cells.

## Contributions

B.E. and M.T.A.S. conceived and designed the experiments. B.E. performed the experiments. B.E., Z.L., M.S.H.J, U.D performed imaging and analysis. B.E., Z.L., M.S.H.J., F.K., M.T.A.S. prepared the manuscript. All authors have read and approved the final manuscript.

## Declaration

The authors declare no competing interests.

## Notes

### Competing Interest Statement

The authors have declared no competing interest.

