## Supplementary figures and images for "A novel method for sensor-based quantification of single/multi-cellular traction dynamics and remodeling in 3D matrices"

### Suppl. Vid. 5

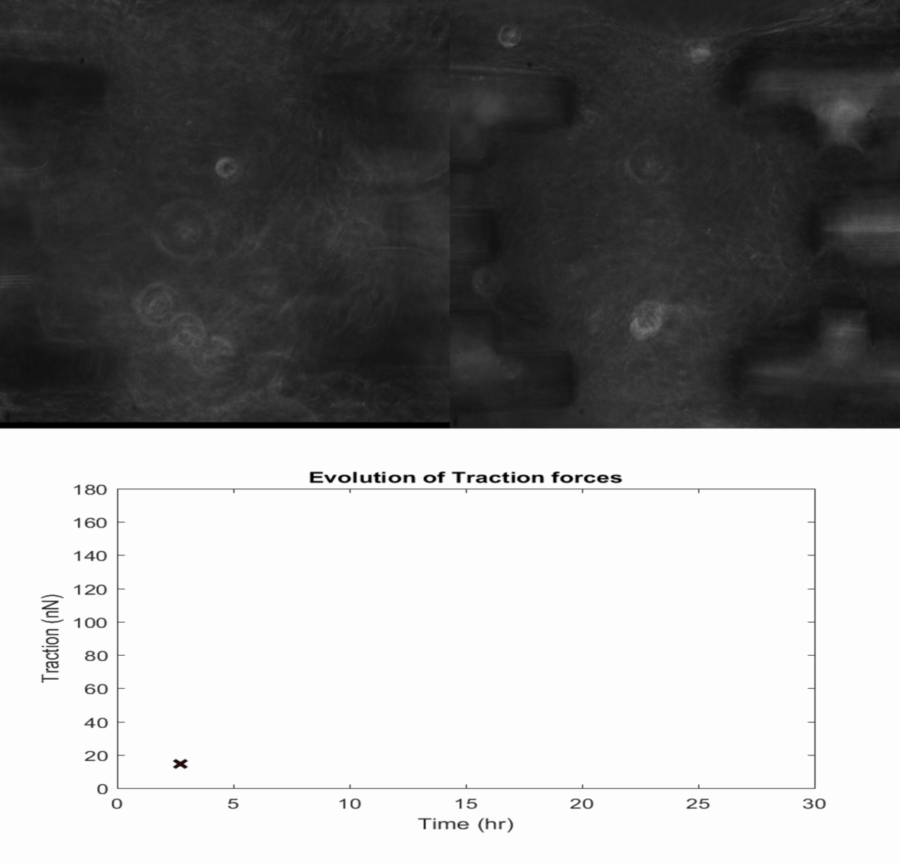
