## Supplementary Document for "A novel method for sensor-based quantification of single/multi-cellular traction dynamics and remodeling in 3D matrices"

### Supplementary Information:

### Calibration and noise analysis

In order to calibrate the sensor springs, we used a glass fiber cantilever beam with known elastic properties as a force calibrator. The procedure is illustrated in Suppl. Fig. 1. Force readout by the sensor spring was found to be accurate within 5% of the actual force applied by the calibrating beam. We also determined the noise level in force data for a collagen specimen without any cells. Contributing factors for noise in the force data can be external vibrations, change of focus and image processing. The Force noise data for 10 hrs of experiment is shown in Suppl. Fig. 2. The data accounts for noise from all possible sources and varies within the range between $\pm 1$ nN.

### Fitting model for Force-strain curves

The non-linear force (F)-strain (ε) relationships of the tension loading segment were fitted to classic constitutive models e.g. Neo-Hookean or Mooney-Rivlin models for hyperelastic polymers [1,2]. For simplicity, we assumed the specimens to be incompressible and hence, for the uniaxial tension experiments, we have the following stress (σ)-stretch (λ) relations:

$$\sigma = 2C_{10}\left(\lambda^2 - {}^{1}/_{\lambda}\right) \quad \text{……………………………………..Neo-Hookean (1)}$$

$$\sigma = 2C_{10}\left(\lambda^2 - {}^{1}/_{\lambda}\right) + 2C_{01}\left(\lambda - {}^{1}/_{\lambda^2}\right) \quad \text{……..….........Mooney-Rivlin (2)}$$

From equations 1 and 2, we can derive the following force (F)-strain (ε) relations respectively:

$$F = 2D_{10}\left((1+\varepsilon)^2 - {}^{1}/_{(1+\varepsilon)}\right) \quad \text{…….………………………………. (3)}$$

$$F = 2D_{10}\left((1+\varepsilon)^2 - {}^{1}/_{(1+\varepsilon)}\right) + 2D_{01}\left((1+\varepsilon) - {}^{1}/_{(1+\varepsilon)^2}\right) \quad \text{..…(4)}$$

All the curves (Fig. 5C-D, Fig. 7D) were fitted to equation 4, except that of FET-CAF05 co-culture specimen (Fig. 7E) which fitted to equation 3. Tangential stiffnesses ($K_t$) of the specimens were determined from the first derivative of F with respect to strain (ε).

### Supplementary Video legends

**Suppl. Vid. 1:** Dispensing of cell-ECM mixture at the tissue location between the grips. For visual clarity, the video was captured without gelatin sacrificial layers that support the sensor at this stage. Movie plays at 10x speed.

**Suppl. Vid. 2:** Single 3T3 cell activities in 3D collagen ECM on the sensor

**Suppl. Vid. 3:** Single CAF05 cell activities in 3D collagen ECM on the sensor

**Suppl. Vid. 4:** Activities and interactions of multiple 3T3 cells in 3D collagen on the sensor

**Suppl. Vid. 5:** Activities and interactions of multiple CAF05 cells in 3D collagen on the sensor

**Suppl. Vid. 6:** Application of compressive and tensile strains in the tissue for (i) stiffness measurement (ii) contraction-stretch assay with cells in 3D ECM

**Suppl. Vid. 7:** 3D reconstruction and surface rendering of the tissue on the sensor

**Suppl. Vid. 8:** Movie with A549 lung cancer model on the sensor

**Suppl. Vid. 9:** Movie with FET CRC tumor model on the sensor

**Suppl. Vid. 10:** Movie with FET and CAF05 co-culture tumor model on the sensor

**Supplementary Figures**

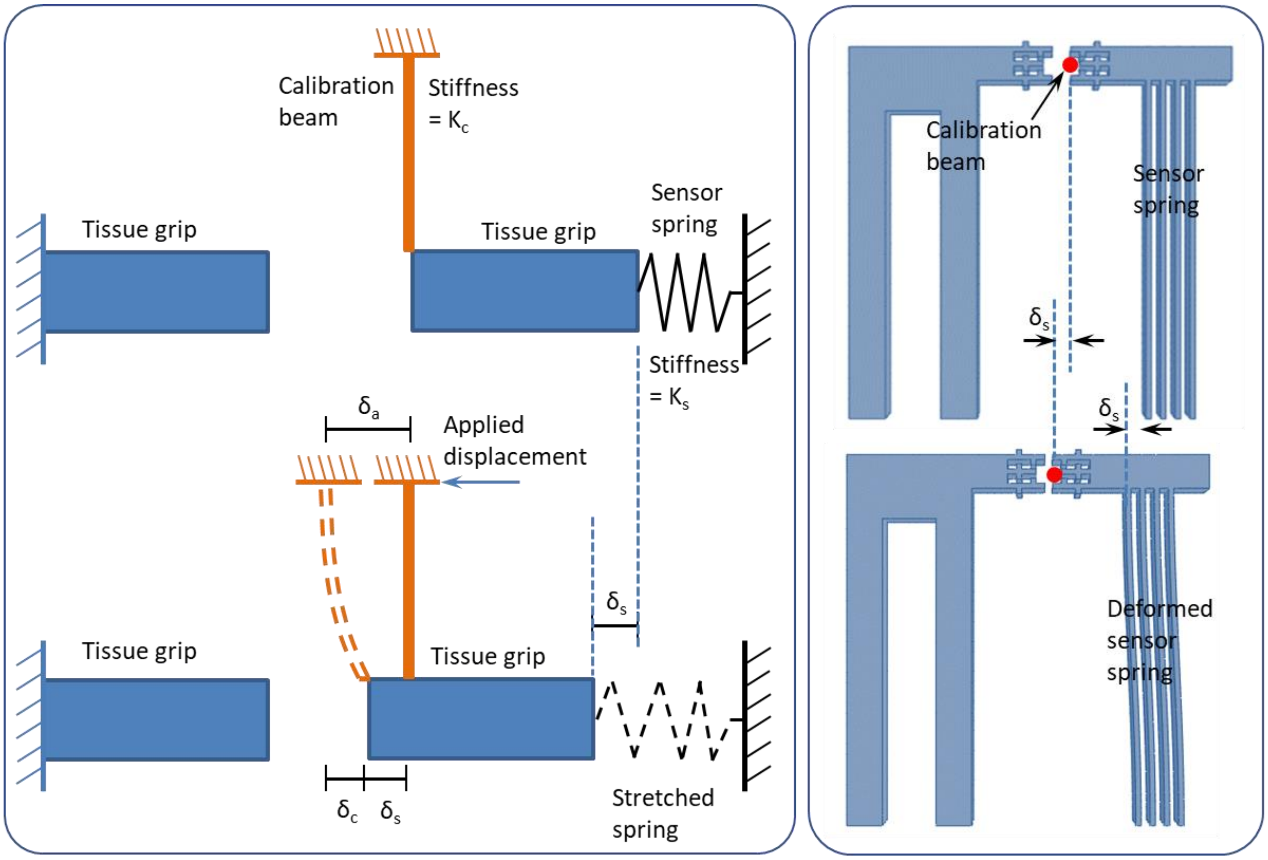


Supplementary Figure 1. Calibration of the sensor spring stiffness . A fiber glass cantilever rod was used as a calibration beam that applied force on the sensing spring. The deflection of the tip of the calibrator ($\delta_c$) and the deformation of the spring ($\delta_s$) are measured using optical microscopy. From known force by the calibrator, the sensor's spring stiffness was determined and compared to the analytic stiffness.

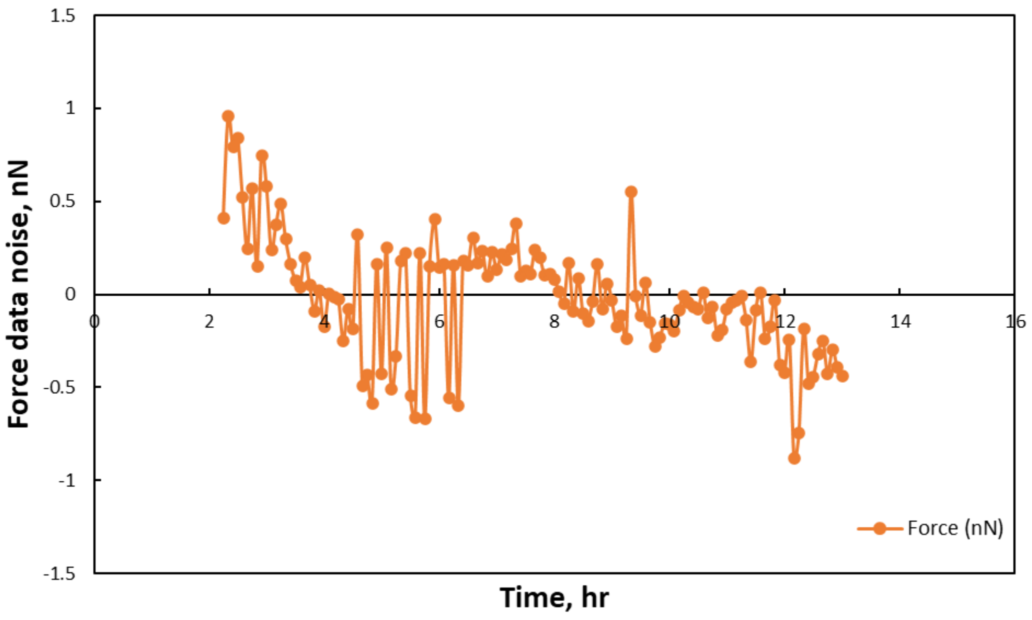


Supplementary Figure 2. Noise data for a specimen without cells.
